# Manifesto of Transparent Mathematical Modeling: From Ecology to General Science

**DOI:** 10.1101/2023.09.19.558506

**Authors:** V. L. Kalmykov, L. V. Kalmykov

## Abstract

Mathematical black box models, which hide the structure and behavior of the subsystems, currently dominate science. Errors and paradoxes, such as the biodiversity paradox and the limiting similarity hypothesis, often arise from subjective interpretations of these hidden mechanisms. To address these problems, we have developed transparent mathematical models of the white box type. Here we justify the hypothesis that transparent mathematical models of the white box type can be built by means of logical deterministic cellular automata whose rules are based on the general theory of the corresponding domain. Using white box models, we were able to directly identify the mechanisms of interspecific competition, test the principle of competitive exclusion and the hypothesis of limiting similarity, resolve the paradox of biodiversity, and formulate for the first time the general principle of competitive coexistence. As a basis for reproducing and further developing the method, we present two transparent mathematical models of an ecosystem with one and two competing species. C++ code for our models provided. Cellular automata thinking can be traced back from ancient cellular board games in the histories of all civilizations. The transparent mathematical modeling opens a rational approach to ensuring the safety, reliability, and trustworthiness of automatic decisions. A shift to transparency in the mathematical modeling paradigm has the potential to revolutionize scientific research and to advance knowledge and technology in a wide variety of domains.

## 1. Introduction

### 1.1. Problems of black box models

Modern mathematical models of complex systems in most cases are based on the methods of continuous mathematics and probability theory. These models belong to the black box type. At the same time, our world consists of clearly discrete systems. The computers we use to simulate these systems are also discrete in nature. Considerable effort has been expended to adapt discrete computers to model discrete objects by methods of continuous mathematics. Non-transparent black-box mathematical models are not able to take into account local conditions of behavior of subsystems of studied objects and are created as a fitting of parameters to the observed phenomenology. Such models are like sketches of observed reality. These mathematical “drawings” can be very precise, but they concern only external properties. As a consequence, conclusions based on opaque models about the internal mechanisms of systems have to be made as indirect subjective interpretations, which generates paradoxes and complicates the understanding of the phenomena under study. A striking attempt to get inside the black boxes of mathematical models of biological systems was N. Rashevsky’s relational biology. Biomathematics, according to Rashevsky, had no way to adequately describe the true integration of the parts of any organic system [1]. Rashevsky began to use the concepts and apparatus of topology for mathematical modeling of the interrelation of subsystems of living organisms [2]. Rashevsky’s student R. Rosen developed this relational approach to living systems, creating the concept of (M,R)-systems [3-5] . Within the framework of this concept, Rosen showed that (M,R)-systems can be interpreted in the language of category theory. Based on the categorical interpretation of (M,R)-systems, Rosen concludes that a living cell can be considered as a set of automata (“general automaton”) and raised the question of the need to create appropriate modeling tools [6]. We believe that a logical deterministic cellular automaton whose rules are based on a general theory of the relevant domain is the very “general automaton” that Rosen wrote about. The work [7] proposes the development of the concept of (M,R)-systems in the sense that an organism is not ‘the complement of a mechanism’: the complement of a mechanism is its ecosystem. In this study, we model an ecosystem as a unity of an organism and all environmental resources necessary for its life. Our Manifesto is a call for the dawn of mathematical biology, caused by the work of N. Rashevsky and R. Rosen [8], to be replaced by a wider use of methods of transparent mathematical modeling of living systems. Rosen sought to ensure that the model accurately reflected the cause-and-effect relationships between the elements of the modeled biosystem [9]. Such a model is of a white box type.

The problem of opaque mathematical models has clearly manifested itself in theoretical ecology with respect to the principle of competitive exclusion. The contradiction of this principle to the observed facts gave rise to what is known as the biodiversity paradox. It turned out to be impossible to check this principle experimentally [10], and the use of mathematical models of the black box type led only to endless fruitless discussions [11-18]. Modern physics in most cases uses opaque mathematical models. A very tense situation has developed in quantum mechanics, in which opacity of used mathematical models has been identified with laws of nature. Principles of quantum mechanics of microcosm are so authoritative today, that they try to extend them to objects of macrocosm, in particular, to ecology. As a consequence, a number of researchers refuse to search for ecological laws, because in their opinion in “an ecological black box opposite ecological concepts according to the principle of ecological uncertainty” borrowed from Heisenberg, “are simultaneously true and false in one and the same measure” and “wave-like combine with each other” [19, 20]. Black box models can reproduce exact phenomenology, but not mechanisms. The mechanisms remain hidden inside the black box. Black box approach can be useful for practical engineering applications. But for the development of science, it is necessary to explore and model the mechanisms. To do this with a black box is, by definition, impossible. This is a feature of the mathematical apparatus. We have shown that mechanisms can be modeled transparently using logical cellular automata. The applied utility of opaque models is not a reason to forget the classical aspirations of science for a transparent rational understanding, including in ecology. 35 years ago, Huston et al. made a vivid call for the use of individual-based methods in ecological modeling, taking into account the local conditions of individuals [21]. According to these authors, this modeling approach would free ecologists from the need to make unreasonable assumptions to model the large-scale behavior of complex systems. We fully agree with this call and believe that to overcome the problems of opaque models it is necessary to create fully transparent white-box type mathematical models. Transparent white box mathematical models are models that are fully structurally and functionally transparent [22, 23]. They allow researchers to easily understand how these models work and the processes that shape them. Transparent white-box mathematical models can bring significant benefits to science and technology. To realize this promising area of research, it is necessary to create models that are as transparent and understandable to researchers as possible, as well as ensuring their high accuracy and efficiency. This will require the cooperation of interdisciplinary teams, including experts in mathematics, computer science, physics, and other sciences.

## 2. On a methodology for transparent mathematical modelling

### 2.1. Definition of a white box model

Transparency of mathematical model of white box type is achieved by a combination of operational and semantic transparency:

*Operational (syntactic) transparency of a mathematical model of a system is a property of the model*, meaning that it is created on the basis of a specific element base while controlling the states and interrelationships of the basic elements within the system. It is “bottom-up” transparency.

*Semantic transparency of mathematical model* is a property of modeling of s-system, meaning the use of explicit concepts, which include the ontology of the highest level. It is “top-down” transparency. The general theory fulfills in transparent model a role of knowledge base of the first principles.

Creation of transparent mathematical model assumes knowledge of algorithms of reproduction of system by logical integration of its elements in complete system. Transparency is closely connected with understanding of mechanism of system functioning. The understanding of mechanism implies possibility to create its transparent mathematical model in imagination or in computer, i.e. transparent model of system is its mechanism.

### 2.2. Our hypothesis

*Completely transparent mathematical models of white box type can be created with the help of deterministic logical cellular automata, which rules are based on the general theory of the subject domain of the system being modeled. It is supposed that the deterministic logical automaton provides operational transparency of mathematical model of the system, and objects and axioms of general theory of the domain of the system being modeled provide its semantic transparency and serve as a knowledge base of first principles*.

Figure 1 shows our classification of mathematical models according to the degree of their transparency, depending on the type of mathematical methods used [24, 25]. It follows from the hypothesis presented by us that a completely transparent model of an object is created by combining the apparatus of cellular automata with the general theory of the corresponding subject area.

**Fig. 1.**
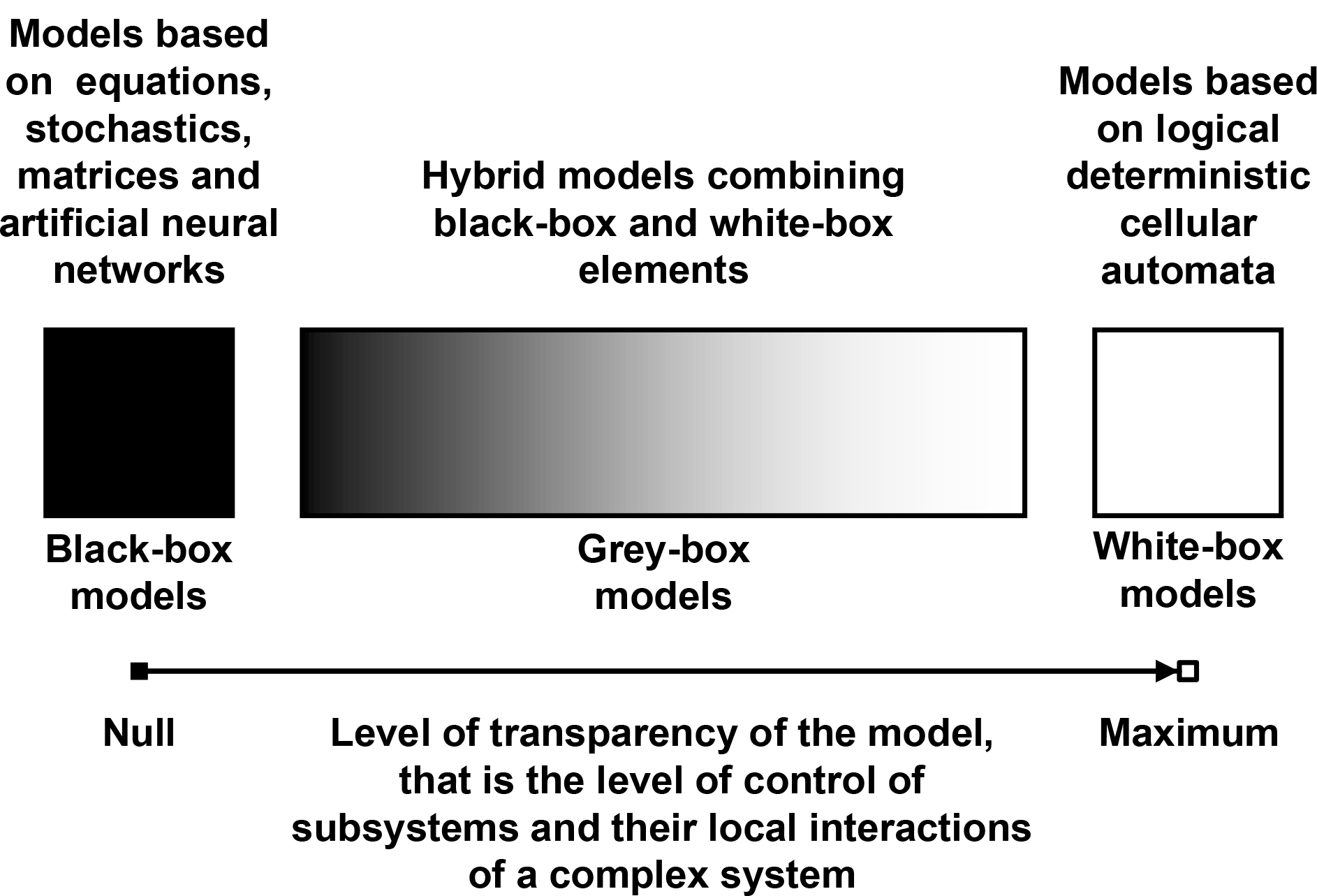
Classification of mathematical models of complex systems according to the degree of their transparency depending on the type of used mathematical methods. This is a schematic representation of black box, grey box and white box type models with indication of the level of control over the cause-effect interactions of subsystems of the integral complex system.

### 2.3. Features and possibilities of cellular automata as the basis of operational transparency of a white box mathematical model

Cellular automata are computational techniques used to model the behavior of complex systems by dividing the system into its basic elements, placed in discrete cells of a periodic lattice. These basic elements interact with their immediate neighbors based on a set of predefined rules. The main advantage of using cellular automata for dynamic simulation is related to their fully discrete nature, which allows for accurate simulation on the computer. Thus, any side effect due to rounding errors is eliminated. This method has found applications in various fields, including physics, theoretical biology and computer science, due to its ability to simulate complex phenomena with simple rules and low computational cost. Cellular automata are effective tools for modeling space, time, and states.

Cellular automata allow creating a transparent model of the system, providing integration of behavior of system elements through their local interactions. “Cellular automata are mathematical idealizations of physical systems in which space and time are discrete and physical quantities take a finite set of discrete values” [26]. A system is modeled by a cellular automaton as an iteratively changing structured set of homogeneous elements. Each element of this set can be in at least two states. States of all elements of this set change stepwise and single-step according to certain logical rules. The rules determine in which state each given element will go at the next step and depend on the state of the element itself and the states of its neighbors given by the local cell-automatic neighborhood. Homogeneous elements coordinated with each other locally in the whole space of the structured set are always and uniquely glued into a coherent global system. The global behavior of the system is generated from local rules of change of states of its microobjects.

Formally a cellular automaton can be defined as five objects:

1. A regular homogeneous lattice of sites (model microobjects);
2. Finite set of possible states of a site;
3. The initial pattern of states of the lattice site.
4. A neighborhood consisting of a site and certain neighboring sites. This set of sites affects every transition of a site to the next state;
5. A transition function to the next state - the rules for iteratively changing the states of each site without exception, which apply to all site of the lattice simultaneously. These rules take into account the neighborhood state of each site. All sites have the same update rules. Iterations of site states model the timeline. The logical causal structure that controls the transition between site states is based on if-then rules.

Each site of a cellular automaton is a finite automaton, and a cellular automaton as a whole is a polyautomaton that organizes the behavior of finite automata located in sites of its lattice.

The cellular automaton implements automatic inference of a multilevel nature. The neighborhood of the cellular automaton indicates the places of applicability of the logical rules of the transition function and represents a spatial pattern. This inextricable link between logic and pattern is an important feature of cellular automata. The logic of the cellular automaton at each iteration is applied simultaneously to simulate changes at all levels of organization of the simulated system. Such multi-level logical inference can be called hyperlogical inference because it causally links changes at different levels of the modeled system into a unified network. The behavior of microobjects of the system being modeled is logically glued into a single coherent macroobject by means of local interactions mediated by the cellular automaton neighborhood. The microlevel is modeled by a lattice site (microecosystem). The mesolevel of local interactions of the microsystem is modeled by the vicinity of cellular automata (mesoecosystem). Macrolevel (the whole ecosystem) is modeled by the whole lattice. It is this modeling, which logically reproduces the integration of subsystems into a coherent system, that allows us to speak of model transparency. Multilevel automatic logical inference performed by cellular-automatic algorithms opens up new opportunities not only for fundamental science but also for the development of new applied methods of transparent artificial intelligence.

### 2.4. Achievement of mathematical model transparency

Deterministic logical cellular automata allow you to control the field of events of the model, starting with the behavior of each individual. We can literally see all the local events on the field. This is a literal understanding of model transparency. But to ensure the physical adequacy of the model, it is not enough just to “draw” the detailed phenomenology of the phenomena under study using mathematical methods. It is necessary to guarantee the absence of inference errors and the fundamental physical correctness of the algorithms underlying the model. The concepts used by the model building algorithms should also be transparent, i.e., explicit. Such fundamental transparency can be achieved if mathematical algorithms are based on first principles, i.e., on basic objects and axioms of the general physical and mathematical theory of the subject area. The use of the basic foundations of the general theory as the first principles of building a model makes it possible to achieve not only mathematical (algorithmic), but also physical adequacy. Physical rigor is achieved using general physical theory as a knowledge base, and deterministic logical cellular automata provide ideal modeling of space and time with full consideration of the role of local conditions in the behavior of each and every individual.

### 2.5. The theory of the subject area as the basis for the semantic transparency of the white box mathematical model

Theory of the subject area accumulates knowledge about objects, properties, relationships, and functional connections in that subject area. This knowledge can be represented as a symbolic mathematical model, which is a formalized description of the objects and their relationships in the subject area. The theory of the subject area serves as a knowledge base for creating a symbolic mathematical model in that subject area. By using this method, a complete understanding of the subject area can be obtained and this understanding can be used to create a symbolic mathematical model.

Diverse transparency is a necessary condition for the development of science and technology in the interests of society. The openness of data, methodologies, and experimental results needs to be complemented by transparency of mathematical models, which is the basis of our understanding of the phenomena being studied. Understanding involves reproducing an object in imagination or on a computer starting from its elemental base. Such an opportunity is provided by scientific rational knowledge. Rational knowledge has a supra-sensory abstract nature, and today the focus of science is on obtaining experimental data and their statistical processing. Abstract results are often considered speculative and are not approved by reviewers. The dominance of positivism in science also does not contribute to the creation of general theories. This problem is one of the limitations of implementing transparent mathematical modeling. The experience of mathematics as a whole has shown that logical operations with symbols corresponding to fully explicit concepts give semantically correct results. To turn partially implicit concepts into explicit ones, it is necessary to construct a general axiomatic theory of the subject area of the modeled object. Such a detailed specification of conceptualization is the construction of the ontology of the subject area [27]. For our research, the abstract theory of living systems [28, 29] served as a general semantic basis. Based on this theory, we formulated a general theory of the ecosystem for studying interspecific competition, which is presented in our article [30] and in earlier works [24, 25, 31] and in this article.

### 2.6. General ecosystem theory

The elementary object of our models is the microecosystem. This is the main ideal object of the theory. The closure of the resources of the microecosystem allows us to consider it as an autonomous object. The closure of ecosystem resources means that all elements within a microecosystem - living organisms, as well as non-living components such as air, water and soil - are interconnected and dependent on each other. Each element of the microecosystem affects its other elements and the microecosystem as a whole. An ecosystem can be considered as an autonomous entity since it has its own characteristics, processes and interactions that exist within it, independent of external factors. The closure of an ecosystem’s resources also means that it can be resilient and able to support the life cycle of a single individual. The autonomous dynamics of this object is characterized by a set of states and a diagram of their successive changes in accordance with extreme physical principles. Extreme physical principles can be characterized as occurring spontaneously according to the second law of thermodynamics. Transitions between the states of the microecosystem are accompanied by the use of free energy. Free energy is used to maintain the life of individuals (the full energy of solar radiation) and to support the processes of restoration of limiting resources during the state of recovery of the microhabitat. In the two-species model, the individual of the dominant species has greater ecological competitiveness and wins direct conflict over resources for its offspring. The individual has the most efficient ability to capture and use free energy and other limiting resources. Following Tansley’s concept [32], we consider individuals with their “special environment with which they form a single physical system”. In the “free” or “occupied” state, the microhabitat contains all the necessary resources for the life of one individual. The flow of high-level solar energy (exergy) ensures the life of an individual in a microhabitat [22, 33, 34]. The flow of low-level energy (anergy) of reduced quality, which is produced during the life of an individual and goes into a cold outer space, also provides conditions for the life of an individual in a microhabitat. A living organism cannot function without the removal of waste heat to the “refrigerator”. Ultimately, the role of such a refrigerator is played by the cold space of space. The working cycle of microecosystem processes includes the occupation of a microhabitat by an individual, the vital activity of an individual, and the restoration of used resources after his death. In our model, vegetative reproduction is carried out at the expense of the resources of the individual’s microhabitat by placing the vegetative rudiment of the offspring in a free microhabitat in the individual’s neighborhood. We consider the plant ecosystem as a “working mechanism” that “supports and restores itself” [35]. To model vegetative reproduction, we use the axiomatics of an excitable environment [36] and the concept of an individual’s microecosystem [25]. The microecosystem is a discrete quantitative unit of the ecosystem and coincides with the previously proposed concept of “environ” [37]. In accordance with this axiomatics of an excitable medium, its three states - rest, excitation and refractoriness successively replace each other. The state of rest corresponds to the free state of the microecosystem, the state of excitation corresponds to the vital activity of the individual, and the refractory state corresponds to the state of regeneration of the microecosystem after the death of the individual. A microhabitat in a regenerating (refractory) microhabitat state cannot be populated by a new individual. Accounting for the state of microhabitat recovery expands the possibilities for modeling ecosystem dynamics, since most ecological models still do not take into account the recovery of resources in plant communities. Previously, the need to take into account the process of restoration of consumed ecosystem resources was discussed by [35] and [38].

### 2.7. Ecosystem Theory Axioms Specifying

1. The space is modeled by a grid with micro-ecosystems/micro-habitats at the sites (Online Supplementary Movies S1 and S2).
2. Time is modeled by successive iterations of state changes of all sites.
3. List of site states (microecosystem/microhabitat):
  - “Free” - can deal with only one descendant of an individual of any species;
  - “Occupied” - the microhabitat is inhabited by an individual and goes into the state of restoration of the conditions and resources of the microhabitat after the death of the individual;
  - “Restoration” - restoration of the conditions and resources of the microecosystem after the death of an individual and disposal of a dead individual of the species. A micro-habitat in the recovery state becomes free or can be occupied immediately after the completion of the recovery state.
4. The cellular automaton neighborhood defines the area of local interactions with each other.
5. The states of the lattice sites (microecosystems/microhabitats) change according to the rules depending on the state of the sites themselves and the states of the sites located in their cellular automaton neighborhood.

One individual can occupy only one microhabitat. The life cycle of an individual, like the state of restoration of a microhabitat, lasts one iteration of the cellular automaton. All states of all sites have the same duration. Each individual of all species consumes the same number of identical resources, i.e. they are identical consumers. Such species are complete competitors. Individuals are immobile at the lattice sites and population waves propagate due to the reproduction of individuals. The neighborhood consists of a site and its given neighboring sites. All sites have the same update rules. The neighborhood models the meso-habitat of the individual and determines the number of possible descendants (fertility) of the individual.

Cellular automata are “bottom-up” models that generate global behavior from local rules [39]. Through bottom-up inference, we were able to ensure that our model mechanistically constructs the ecosystem from the bottom up at each iteration, from the local conditions of each individual microhabitat and each individual species to the ecosystem as a whole. The rules of cellular automata are implemented at three levels of ecosystem organization. It is this multilevel modeling that makes it possible to speak about the transparency of this method. The microlevel is modeled by a lattice site (microecosystem). The mesolevel of local interactions of a microecosystem is modeled by the neighborhood of cellular automata (mesoecosystem). The macro level (the entire ecosystem) is modeled by the entire grid. The logical rules of the model include both “part-whole” relationships and “cause-effect” relationships. The whole is the entire cellular automaton, and the parts are the cells and their neighborhoods.

### 2.8. On the transparent cellular automata thinking

The use of a cellular automata approach for modeling was implemented in ancient board games. These games developed logical and visual thinking skills in the field of strategy and tactics. Chess is the most well-known logical game, modeling the battle of two armies on a field. The oldest known strategic games on a grid are Senet and the Royal Game of Ur. Senet was invented in Ancient Egypt around 5500 years ago. The Royal Game of Ur was invented in Sumer at least 4400 years ago. In Ancient Greece, people played Petteia on boards of 6x6, 7x7, and 8x8. Many vase paintings have survived depicting scenes where Achilles and Ajax play board games, presumably Petteia. In Ancient Rome, the game of Latrunculi (“little soldiers”) was popular, with many variations in rules and board size. Go is a logical strategic board game that originated in Ancient China, estimated to be between 2 and 5 thousand years ago. The game of Patolli has a long history and wide geography in pre-Columbian America: it was played by the Teotihuacans (about 200 BC - 650 AD), Toltecs (750-1000 AD), Chichen Itza residents (1100-1300 AD), and Aztecs (1168-1521 AD). Ancient Mayans played a variety of Patolli.

We believe that in order to create a board game on a grid, it is first necessary to create a theory. Creating a theory means defining the basic objects of the theory (pieces, board) and the axioms of the behavior of these basic objects (game rules). This classical rational approach to modeling emerged spontaneously and independently in different civilizations. The prevalence of logical board games on a grid in all advanced civilizations testifies to their unique effectiveness in modeling complex systems and the exciting interest they have aroused in players, providing complete visibility and transparency of all events on the board. We consider these games to be cellular automata, as they possess similar characteristics: (1) the board consists of ordered cells, (2) each cell can be in a finite number of known states, (3) pieces have specific local neighborhoods, (3) there is an initial pattern of the board (4) transitions between cell states are realized iteratively according to certain logical rules. Given the prevalence of cellular automata stereotypes in board games of different civilizations, it can be assumed that cellular automata thinking is natural for humans. Perhaps our thinking uses cellular automata modeling of the world in which we live. At the same time, human imagination is not sufficient to involve all the pieces of opposing sides in every iteration of the game. As a result, the game is played by alternating individual moves of individual pieces. Computers have opened up the possibility of modeling changes in the states of all cells on the board in each iteration, which has increased the realism of the models, as many processes in nature occur simultaneously. The spontaneous cellular automata nature of board games on a grid is a unique confirmation of the idea that the Universe is a cellular automaton, as these games are very effective models of reality.

### 2.9. Is our universe a cellular automaton?

The deterministic dynamics of cellular automata is considered a candidate for the fundamental laws of physics, including quantum theory [40, 41]. The effectiveness of cellular automata modeling may be related to the fact that the fundamental laws of nature have a cellular automaton nature, and the universe itself is a cellular automaton. Conrad Zuse previously proposed such a hypothesis in his book “Computing Space” [42]. Zuse calls for an “automaton way of thinking” in physics, which implies that a physical model is considered from the point of view of a sequence of states following each other according to pre-defined rules. Following Zuse, Edward Fredkin believed that the universe is a computer program that evolves on a grid according to a simple rule [43]. If we could only determine the correct type of grid and the correct rule, says Fredkin, we could model the universe and all of physics. Fredkin insists that the rule governing the cellular automaton of the universe must be simple - just a few lines of clean and elegant code. Continuing the ideas of K. Zuse and E. Fredkin, Stephen Wolfram attempted to find the digital cellular automaton code of the universe. Twenty years ago, he tried to find this code by enumerating rules and patterns of states of sites in the lattice of simple cellular automata [44].

Today, Wolfram has supplemented these searches by enumeration of the rules of transformation graphs of sets of relationships of identical but labeled discrete elements [45, 46]. The approach is similar to cellular automaton modeling represented by state transition graphs, but with the introduction of the maximum degree of arbitrariness by relaxing the cellular automaton formalism. As a result of this arbitrariness, the possibilities of adjusting the resulting models to physical phenomenology are expanding in the hope of finding the same common code of the Universe. The code of the Universe has not yet been found, and Wolfram’s project to search for a fundamental theory of physics has been criticized [47]. It is unlikely to find the rules for the creation of the universe just by combining numbers. Even if cellular automata and graphs are used. We share the idea that our world is based on cellular automaton logic, but we believe that a more meaningful search for an appropriate fundamental theory is needed. It is necessary to find the first principles (objects and axioms) of the general physical theory that underlie the rules of the cellular automaton of our Universe.

## 3. The examples of the implementation of transparent mathematical models of the white box type

### 3.1. The white box models of ecosystem with one and two competing species

Textbooks on mathematical modeling often present classical models of population dynamics. These are the classical models of Malthus, Verhulst and Lotka-Volterra. They model population growth of one and two species. These are models based on differential equations, which are black boxes, since they do not model the internal structure of the population and the mechanisms of local interactions of individuals. Our research discusses how and why the use of opaque models in ecology led to a number of serious fundamental problems and paradoxes, some of which we managed to overcome with the help of transparent mathematical modeling [24, 25, 30, 31, 48-50]. Here we provide the code and description of two ecosystem models with one and two competing species that were used in our studies. In contrast to the models of Malthus, Verhulst and Lotka-Volterra, they are of the “white box” type and allow one to visually explore local mechanisms. This ensures a rigorous interpretation of the results.

Both models are logical deterministic cellular automata that implement automatic logical inference at three levels of organization of a complex system. These are examples of the implementation of transparent artificial intelligence based on transparent mathematical modeling. Both models are good examples of white box models. These examples support our hypothesis that logical deterministic cellular automata make it possible to model complex systems in a visually transparent way, creating white-box models. The results obtained by solving old scientific paradoxes are a breakthrough in a fundamental scientific sense. At the same time, this is a precedent for the implementation of innovative transparent artificial intelligence technology based on white-box modeling. The program code given in Supplementary Information to this article allows researchers to use these programs as a universal framework for further development of transparent mathematical modeling. The program code is written in C++ and tested in Microsoft Visual C++ Studio 2015. Comments in the code are highlighted in green. These programs implement computer experiments. The code allows you to set the size of the habitat, the initial location of individuals in the habitat. The cellular automaton lattice (microhabitat) consists of 25x25 sites (microhabitats). For large lattice sizes, for clarity, you need to increase the size of the application window and reduce the font in the console properties. The initial pattern of the ecosystem consists of free space inhabited in the first model by one individual of Species 1 and in the second model by two single individuals of two competing species. Under given initial conditions, the programs show classical examples of population growth and demonstrative local mechanisms of population dynamics. It looks like the propagation, collision and interaction of population waves of one and two species in a limited environment with renewable resources. Therefore, these population waves are autowaves. That is, from a physical point of view, Model 1 demonstrates the propagation of one type of autowaves in a limited environment with renewable resources (Online Supplementary Movie S1). And Model 2 demonstrates the propagation and interaction of two different types of autowaves in a limited environment with renewable resources (Online Supplementary Movie S2). To avoid edge effects, both models implement periodic boundary conditions that close the lattice of the cellular automaton into a torus. From an ecological point of view, the free habitat is colonized by individuals of one and two competing species.

### 3.2. From the metaphors of the game of life to our precedents for the implementation of completely transparent models

The computer cellular automaton game “Life” (Game of Life) by John Conway [51] has gained great popularity. The player arbitrarily creates a pattern of “live” (shaded) and “dead” (unshaded) cells on a two-dimensional field, and then only observes the evolution of the patterns. If the number of “living” neighboring cells is 2 or 3 out of 8, then at the next iteration the cell “survives”, if not, then it dies. A “dead” cell “comes to life” at the next iteration if only it is surrounded by three “live” cells. The player hopes to get interesting dynamics during the evolution of the machine. Of course, this is not a model of a specific object of nature, but a very distant metaphor that can be attributed to abstract art. But this metaphorical model has analogies in many disciplines and allows you to reproduce dynamic processes that were not amenable to mathematical modeling before. For us, this model is important because, having taken its program [52] as a basis, we have replaced its rules taking into account the objects and axioms of the general theory of an ecosystem with competing species. The ecological model of resource competition implemented by us has a completely transparent nature, unlike all available solutions [24, 25, 30] and allowed us to solve a number of problems of theoretical ecology:

1. For the first time, completely transparent individual-based mechanisms for the formation of classical population growth curves, including the double S-shaped curve, were created and studied [48].
2. For the first time, a discrete model of population catastrophes was created; it was demonstrated that with an increase in the recovery time of ecosystem resources, a catastrophic death of the population occurs [49].
3. Mechanisms have been found for the indefinitely long coexistence of full resource competitors under conditions under which coexistence was previously considered impossible. This solved the paradox of biodiversity and opened up new ways to conserve biodiversity [24, 25, 50].
4. For the first time, the competitive exclusion principle was rigorously tested and reformulated, and a generalized formulation of the competitive exclusion principle for an arbitrary number of resource competitors was given [24, 25, 50].
5. Two contradictory hypotheses of limiting similarity and limiting difference of competing species were tested; the hypothesis of limiting similarity was rejected [30].
6. For the first time, the general principle of competitive coexistence was formulated for an arbitrary number of resource competitors [25, 50].

These results confirm the legitimacy of our hypothesis that a completely transparent model of a complex system can be created by connecting the apparatus of cellular automata with the axioms of the general theory of the relevant subject area. The obtained results also confirm that *a deterministic logical cellular automaton is able to provide the operational transparency of the mathematical model of the system, and the objects and axioms of the general theory of the subject area of the modeled system are able to provide its semantic transparency*.

The details of our transparent mathematical modeling method are published in our papers and briefly presented in the appendix to this paper. The Supplementary Information contains program codes for one-species and two-species models.

## 4. Conclusion and perspectives

The presented study demonstrates the possibility of transparent multilevel modeling of complex systems as a result of combining the axiomatic theory of ecosystems with the formalism of a logical cellular automaton. We consider the general theory of the ecosystem as a theory that is of fundamental importance not only for population ecology, but also for mathematical modeling in general. The fact is that any active structure can be considered as an agent with a certain degree of autonomy and the ability to interact with its environment. From this point of view, any active structure can be considered as an agent with its own ecosystem. A fundamentally important point is that it is not the active structure itself that acts as a completely autonomous agent, but its integral complex with its inherent habitat. In our models, an individual (an individual of a species capable of vegetative reproduction) acts as an active agent, and the ecosystem of an individual is the unity of an individual with its microhabitat. The integral complex of an individual and his micro-habitat is a micro-ecosystem (an elementary basic object of the theory), which has true autonomy. This generalizing point of view in application in our general theory of the ecosystem can be applied in various fields, including biology, sociology, economics and artificial intelligence.

Our approach as a whole opens up great prospects for various theoretical and applied fields. The paper is a reflection on the precedents of using a fully transparent method for modeling the behavior of multiparticle and multilevel systems. This work is written as a harbinger of the coming spring of transparent mathematical modeling based on symbolic rational knowledge, on clearly formulated algorithms, on logic, on a general picture of the world. The implementation of the proposed program requires students to become more familiar with both cellular automata thinking and the rational scientific method. We assume that transparent and non-transparent methods of mathematical modeling will harmoniously complement each other in hybrid solutions. A shift in the paradigms of mathematical modeling towards completely transparent solutions has the potential to revolutionize scientific research and expand our knowledge in many areas.

## Supporting information

A white box model of colonization of free ecosystem by one species (Movie S1)

A white box model of colonization of free ecosystem with resource competition between two species (Movie S2)

## Data availability statement

The authors declare that all data used to create the Online Supplementary Movies S1 and S2 related to this article are contained in the movies themselves. Movies reflect both graphic information and digital data from computer experiments.

## Supplementary materials

The supplementary materials include Supplementary Information and Supplementary Online Movies S1 and S2.

## Author contributions

All authors contributed to the study conception. The manuscript was written by both authors. Programs and movies of experiments were created by L.V. Kalmykov. All authors read and approved the final manuscript.

## Acknowledgements

We thank Lev Alexandrovich Naumov for permission to use his source code for John Conway’s Game of Life as a prototype.

