## Supplementary Information for "Manifesto of Transparent Mathematical Modeling: From Ecology to General Science"

V. L. Kalmykov[1,*], L. V. Kalmykov[2]

[1] Institute of Cell Biophysics of the Russian Academy of Sciences, 3 Institutskaya st., Pushchino, Moscow region, 142290, Russian Federation
[2] Institute of Theoretical and Experimental Biophysics of the Russian Academy of Sciences, 3 Institutskaya st., Pushchino, Moscow region, 142290, Russian Federation

*

### Description of the program code of transparent ecosystem models with one and two competing species

Here we provide a description and code of our models. The biological prototype of the models is the vegetative propagation of rhizome-loose-shrub lawn grasses (Poa pratensis L., Festuca rubra L. ssp. Rubra and their varietal variations) in a homogeneous limited habitat. Vegetative reproduction excludes the occurrence of genetic diversity of individuals of the species. For us, the genetic homogeneity of individuals of the same species was an important condition in testing the formulations of the competitive exclusion principle and the competing hypotheses of limiting similarity and limiting difference. Rhizomes of lawn grasses distribute vegetative descendants in the form of two-dimensional network structures. One individual corresponds to one single shoot. A single grass shoot is relatively autonomous, germinates from a bud at the end of a rhizome and is able to reproduce. Rhizomes are horizontal creeping underground rhizome shoots, with the help of which plants reproduce vegetatively. Unlike roots, rhizomes at the end have buds with rudimentary scaly leaves and nutrient reserves for the initial development of single shoots with roots and leaves. One single shoot in the presented model can have a maximum of six rhizome processes within a hexagonal neighborhood. The hexagonal neighborhood most naturally corresponds to the dense packing of round microhabitats of single grass shoots. By changing the type and rank of the neighborhood, one can simulate various features of reproduction [1, 2].

The program code with the description allows using it as a basic basis for the further development of transparent mathematical modeling. As an example, we took a field of 25x25 cells. The two-dimensional hexagonal lattice is closed on the torus by periodic boundary conditions in order to avoid boundary effects. We use a hexagonal grid because it most naturally implements the densest packing of microhabitats. The hexagonal neighborhood makes it possible to model potentially aggressive vegetative propagation of plants, when the offspring of an individual can occupy all the nearest microhabitats (Supplementary Fig. 1a).

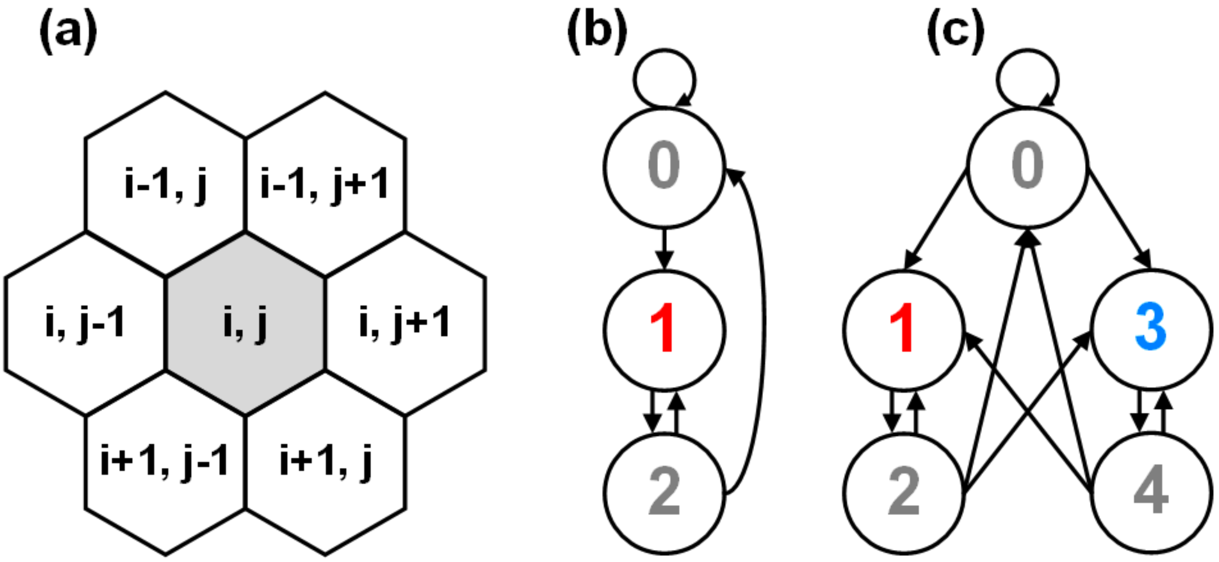


**Supplementary Figure 1. Rules of a fully transparent logical model of an ecosystem with one species and two closely competing species.** (a), a hexagonal neighborhood. The indexes of the elements of the array i and j are given as integers. (b), A diagram of transitions between cell states of a cellular automaton field in a single-species ecosystem model. (c), A diagram of transitions between the cell states of a cellular automaton field in an ecosystem model with two closely competing species. The list of possible states in the models: "0" — the microhabitat is free; "1" - the microhabitat is inhabited by an individual of Species 1; "2" — a microhabitat in a state of regeneration after the death of an individual of Species 1; "3" — a microhabitat inhabited by an individual of Species 2; "4" - a micro-habitat in a state of regeneration after the death of an individual of Species 2. The logical rules that determine the conditions for transitions between the states of the lattice site of the cellular automaton are visually represented by the transition diagram (Supplementary Figure b, c) and in the program code with comments. The description of the variables in the programs of both models is given in Supplementary Table 1.

**Supplementary Table 1. Description of variables in the programs for Model 1 and Model 2**

| Integer variable | Описание |
|---|---|
| n | The number of rows in the arrays Array[n][m] and Array_temp[n][m]. These are arrays for storing the current and calculated states of lattice sites. |
| m | The number of columns in arrays Array[n][m] Array_temp[n][m].<br>Total number of grid sites: n * m |
| s | The state in which the central site of the neighborhood of the cellular automaton is located. It is defined by the array element with indexes (i, j). |
| s1 | The state of the site of the neighborhood of the cellular automaton. It is defined by the array element with indexes (i-1, j). |
| s2 | The state of the site of the neighborhood of the cellular automaton. It is defined by the array element with indexes (i-1, j+1). |
| s3 | The state of the site of the neighborhood of the cellular automaton. It is defined by the array element with indexes (i, j+1). |
| s4 | The state of the site of the neighborhood of the cellular automaton. It is defined by the array element with indexes (i+1, j). |
| s5 | The state of the site of the neighborhood of the cellular automaton. It is defined by the array element with indexes (i+1, j-1). |

| | |
|---|---|
| s6 | The state of the site of the neighborhood of the cellular automaton. It is defined by the array element with indexes (i, j-1). |

### Description of the first model: an ecosystem with one species

Model 1 simulates an ecosystem with a single species in conditions of limited resources. A visual mechanism of propagation of population autowaves in a limited homogeneous environment as a result of reproduction of individuals is shown. In this case, the classical S-type population curve is realized. Previously, this model was used by us to study the conditions of the catastrophic death of a single-species population. Unlike the Verhulst model, which is a black box, this model is a white box, i.e., a complete transparent model of the ecosystem [3, 4].

A description of all possible states of each space element for Model 1 is given in Supplementary Table 2.

**Supplementary Table 2. Description of possible states of the lattice site in Model 1**

| Symbol | Description |
|---|---|
| 0 | This is a state of free micro-habitat that contains resources and can be populated. |
| 1 | This is a state of micro-habitat that is inhabited by an individual of Species 1. |
| 2 | This is a state of micro-habitat regeneration with the restoration of all resources and conditions after the death of an individual of Species 1. |

Rules for transitions between site states (microecosystems) in the ecosystem model with one species (Supplementary Figs. 1a, b):

0 → 0, the microhabitat remains free if there is not a single living individual in the neighborhood;

0 → 1, if there is at least one individual of Species 1 in the vicinity of the cellular automaton, then the microhabitat is populated by an individual of Species 1;

1 → 2, after the death of an individual of Species 1, his micro-habitat goes into a state of restoration;

2 → 0, after the recovery state, the micro-habitat will be free if there is not a single living individual in the neighborhood;

2 → 1, if there is at least one individual of Species 1 in the vicinity of the cellular automaton, then the microhabitat after the restoration of resources is populated by an individual of Species 1.

### Program code of the first model: ecosystem with one species

```
#include "stdafx.h"
#include <iostream>
#include <conio.h>
#include <windows.h>

using namespace std;

// Grid size (ecosystem) 25x25 sites (micro-habitats)
const int n = 25; // The number of rows in the array
const int m = 25; // The number of columns in the array

int s[n][m]; // Array for storing the current states of the lattice sites
int s_temp[n][m]; // Array for storing calculated states of lattice sites

// Periodic boundary conditions closing the lattice in the torus
```

```
int Torus_n(int i)
{
	if (i<0) return i + n;
	else return i % n;
}

int Torus_m(int j)
{
	if (j<0) return j + m;
	else return j % m;
}

/* The function of transitions between the states of the lattice site of the cellular
automaton. The function contains the rules of automatic logical output.*/
int f1(int s, int s1, int s2, int s3, int s4, int s5, int s6)
{

/* Free micro-habitat and micro-habitat after the regeneration state is occupied by the
descendant of an individual, if there is at least one individual in the neighborhood */
if (((s1 == 1) || (s2 == 1) || (s3 == 1) || (s4 == 1) || (s5 == 1) || (s6 == 1)) &&
		((s == 0) || (s == 2)))
		return 1;

/* After the death of an individual, the micro-habitat goes into a state of resource
regeneration */
if (s == 1) return 2;

/* After the regeneration state, the micro-habitat becomes free if there is not a single
individual in the neighborhood */

if (s == 2) return 0;

/* The micro-habitat remains free if there is not a single individual in the neighborhood
*/
if (s == 0) return 0;
}


int main()
{
	system("Color F0"); // Field color - white
	int iteration = 0; // Initial iteration
	setlocale(LC_ALL, ""); // Russian font

	// Initial filling of arrays
	for (int i = 0; i<n; i++)
		for (int j = 0; j<m; j++)
		{
			s[i][j] = 0;
			s_temp[i][j] = 0;
		}

	s[12][12] = 1; // Initial placement of an individual in a habitat

	for (;;)
	{
// Calculating the number of individuals in a habitat
		int a = 0;
		for (int i = 0; i<n; i++)
			for (int j = 0; j<m; j++)
			{
				if (s[i][j] == 1) a++;
			}

// Console visualization
```

```
            cout << "\n";
            for (int i = 0; i<n; i++)
            {
                  for (int k = 0; k <= i; k++)
                        cout << " ";
                  for (int j = 0; j<m; j++)

                  {
                        if (s[i][j] == 0)
                        {
                              HANDLE hOut;
                              hOut = GetStdHandle(STD_OUTPUT_HANDLE);
                              SetConsoleTextAttribute(hOut, BACKGROUND_RED |
BACKGROUND_GREEN | BACKGROUND_BLUE | BACKGROUND_INTENSITY | 7);
                              cout << " 0";
                        }

                        if (s[i][j] == 1)
                        {
                              HANDLE hOut;
                              hOut = GetStdHandle(STD_OUTPUT_HANDLE);
                              SetConsoleTextAttribute(hOut, BACKGROUND_RED |
BACKGROUND_GREEN | BACKGROUND_BLUE | BACKGROUND_INTENSITY | FOREGROUND_RED |
FOREGROUND_INTENSITY | 0);
                              cout << " 1";
                        }

                        if (s[i][j] == 2)
                        {
                              HANDLE hOut;
                              hOut = GetStdHandle(STD_OUTPUT_HANDLE);
                              SetConsoleTextAttribute(hOut, BACKGROUND_RED |
BACKGROUND_GREEN | BACKGROUND_BLUE | BACKGROUND_INTENSITY | 7);
                              cout << " 2";
                        }
                  }     cout << "\n";
            }

// Data output
            HANDLE hOut;
            hOut = GetStdHandle(STD_OUTPUT_HANDLE);
            SetConsoleTextAttribute(hOut, BACKGROUND_RED | BACKGROUND_GREEN |
BACKGROUND_BLUE | BACKGROUND_INTENSITY);

            cout << " Number of iterations: " << iteration << "\n";
            cout << " The number of individuals of the Species 1: " << a << "\n";

// Calculating the states of the lattice sites in the next iteration
            for (int i = 0; i<n; i++)
            for (int j = 0; j<m; j++)

/* Calculating the state of the lattice site in the next iteration. The arguments of the
f1 function are the state of the central site and the states of sites from its
neighborhood. */
            s_temp[i][j] = f1(s[Torus_n(i)][Torus_m(j)],
                              s[Torus_n(i - 1)][Torus_m(j)],
                              s[Torus_n(i - 1)][Torus_m(j + 1)],
                              s[Torus_n(i)][Torus_m(j + 1)],
                              s[Torus_n(i + 1)][Torus_m(j)],
                              s[Torus_n(i + 1)][Torus_m(j - 1)],
                              s[Torus_n(i)][Torus_m(j - 1)]);

/* Transfer of new states of lattice sites to an array of current states of lattice
sites. */
            for (int i = 0; i<n; i++)
```

```
            for (int j = 0; j<m; j++)
            s[i][j] = s_temp[i][j];

            _getch(); // Pause
            iteration++; // Iteration counter
            system("cls"); // Clearing the screen
        }
        return 0;
}
```

### Description of the second model: an ecosystem with two competing species

Model 2 simulates an ecosystem with two competing species in the same way under conditions of limited resources. A visual mechanism of propagation and collision of population autowaves of two Species in a limited homogeneous environment as a result of the reproduction of individuals competing for microhabitats free for settlement is shown. At the same time, the familiar population curves from the classical experiment of G. F. Gause with the competition of two close species of paramecia appear on the graphs of the number of populations [5]. In this model case, the dominant Species 1 completely displaces the recessive Species 2. Previously, this model was used by us to test the well-known formulations of the principle of contingent exclusion. For the first time, deterministic mechanisms of coexistence of full resource competitors were found, which led to the creation of a strict formulation of the principle of competitive exclusion and the solution of the bio-diversity paradox. In the future, this model was used to test the hypotheses of limiting similarity and limiting difference. Unlike the competition model of two similar Species of Lotka-Volterra, which is a black box, this model is a white box, i.e., the most complete transparent model [2, 6].
A description of all possible states of each space element for Model 2 is given in Supplementary Table 3.

**Supplementary Table 3. Description of possible lattice site states in Model 2**

| Symbol | Description |
|---|---|
| 0 | This is a state of free micro-habitat that contains resources and can be populated. |
| 1 | This is a state of microhabitat that is inhabited by an individual of Species 1. |
| 2 | This is a state of micro-habitat regeneration with the restoration of all resources and conditions after the death of an individual of Species 1. |
| 3 | This is a state of micro-habitat that is inhabited by an individual of Species 2. |
| 4 | This is a state of micro-habitat regeneration with the restoration of all resources and conditions after the death of an individual of Species 2. |

Rules for transitions between site states (microecosystems) in an ecosystem model with competition for resources between two species (Supplementary Figs. 1a, c):

0 → 0, microhabitat remains free if there is not a single living individual in the neighborhood;

0 → 1, if there is at least one individual of Species 1 in the neighborhood of the cellular automaton, then the microhabitat is populated by an individual of Species 1;

0 → 3, if there is no individual of Species 1 in the neighborhood of the Species 1, then the microhabitat is populated by an individual of Species 2;

1 → 2, after the death of an individual of Species 1, his micro-habitat goes into a state of regeneration;

2 → 0, after the recovery state, the micro-habitat will be free if there is not a single living individual in the neighborhood;

2 → 1, if there is at least one individual of Species 1 in the vicinity of the cellular automaton, then the microhabitat after the restoration of resources is populated by an individual of Species 1;

2 → 3, if there is not a single individual of Species 1 in the vicinity of the cellular automaton, but there is at least one individual of Species 2, then the microhabitat after the restoration of resources is populated by an individual of Species 2;

3 → 4, after the death of an individual of Species 2, his micro-habitat goes into a state of regeneration;

4 → 0, after the recovery state, the micro-habitat will be free if there is not a single living individual in its vicinity;

4 → 1, if there is at least one individual of Species 1 in the vicinity of the cellular automaton, then the microhabitat after the restoration of resources is populated by an individual of Species 1;

4 → 3, if there is not a single individual of Species 1 in the vicinity of the cellular automaton, but there is at least one individual of Species 2, then the microhabitat after the restoration of resources is populated by an individual of Species 2.

### Program code of the second model: ecosystem with two species

```
#include "stdafx.h"
#include <iostream>
#include <conio.h>
#include <windows.h>

using namespace std;

// Grid size (ecosystem) 25x25 sites (micro-habitats)
const int n = 25; // The number of rows in the array
const int m = 25; // The number of columns in the array

int s[n][m]; // Array for storing the current states of the lattice sites
int s_temp[n][m]; // Array for storing calculated states of lattice sites

// Periodic boundary conditions closing the lattice in the torus
int Torus_n(int i)
{
	if (i<0) return i + n;
	else return i % n;
}

int Torus_m(int j)
{
	if (j<0) return j + m;
	else return j % m;
}

/* The function of transitions between the states of the lattice site of the cellular
automaton. The function contains rules for automatic logical inference. */
int f1(int s, int s1, int s2, int s3, int s4, int s5, int s6)
{
```

```
/* Free micro-habitat and micro-habitat after the regeneration state is occupied by a
descendant of an individual of Species 1, if there is at least one individual of Species
1 in the neighborhood */
        if (((s1 == 1) || (s2 == 1) || (s3 == 1) || (s4 == 1) || (s5 == 1) || (s6 == 1))
&&      ((s == 0) || (s == 2) || (s == 4)))
        return 1;

/* Free micro-habitat and micro-habitat after the regeneration state is occupied by the
descendant of an individual of Species 2, if there is at least one individual of Species
2 in the neighborhood */
        if (((s1 == 3) || (s2 == 3) || (s3 == 3) || (s4 == 3) || (s5 == 3) || (s6 == 3))
&&      ((s == 0) || (s == 2) || (s == 4)))
        return 3;

        if (s == 1) return 2; // death of an individual of Species 1
        if (s == 2) return 0; // Micro-habitat becomes free
        if (s == 3) return 4; // Death of an individual of Species 2
        if (s == 4) return 0; // Micro-habitat becomes free
        if (s == 0) return 0; // Micro-habitat remains free
}


int main()
{
        system("Color F0"); // Field color - white
        int iteration = 0; // Initial iteration
        setlocale(LC_ALL, ""); // Adding Fonts

        // Initial filling of arrays
        for (int i = 0; i<n; i++)
        for (int j = 0; j<m; j++)
                {
                        s[i][j] = 0;
                        s_temp[i][j] = 0;
                }

        s[10][10] = 1; // Initial placement of an individual of Species 1
        s[14][14] = 3; // Initial placement of an individual of Species 2

        for (;;)
        { // Calculating the number of individuals on the field
                int a = 0;
                int b = 0;
                for (int i = 0; i<n; i++)
                        for (int j = 0; j<m; j++)
                        {
                                if (s[i][j] == 1) a++; // Number of individuals of Species 1
                                if (s[i][j] == 3) b++; // Number of individuals of Species 2

                        }

// Console visualization
                cout << "\n";
                for (int i = 0; i<n; i++)
                {
                        for (int k = 0; k <= i; k++)
                                cout << " ";
                        for (int j = 0; j<m; j++)

                        {
                                if (s[i][j] == 0)
                                {
                                        HANDLE hOut;
                                        hOut = GetStdHandle(STD_OUTPUT_HANDLE);
```

```
                                SetConsoleTextAttribute(hOut, BACKGROUND_RED |
BACKGROUND_GREEN | BACKGROUND_BLUE | BACKGROUND_INTENSITY | 7);
                                cout << " 0";
                        }

                        if (s[i][j] == 1)
                        {
                                HANDLE hOut;
                                hOut = GetStdHandle(STD_OUTPUT_HANDLE);
                                SetConsoleTextAttribute(hOut, BACKGROUND_RED |
BACKGROUND_GREEN | BACKGROUND_BLUE | BACKGROUND_INTENSITY | FOREGROUND_RED |
FOREGROUND_INTENSITY | 0);
                                cout << " 1";
                        }

                        if (s[i][j] == 2)
                        {
                                HANDLE hOut;
                                hOut = GetStdHandle(STD_OUTPUT_HANDLE);
                                SetConsoleTextAttribute(hOut, BACKGROUND_RED |
BACKGROUND_GREEN | BACKGROUND_BLUE | BACKGROUND_INTENSITY | 7);
                                cout << " 2";
                        }

                        if (s[i][j] == 3)
                        {
                                HANDLE hOut;
                                hOut = GetStdHandle(STD_OUTPUT_HANDLE);
                                SetConsoleTextAttribute(hOut, BACKGROUND_RED |
BACKGROUND_GREEN | BACKGROUND_BLUE | BACKGROUND_INTENSITY | FOREGROUND_BLUE |
FOREGROUND_INTENSITY | 0);
                                cout << " 3";
                        }

                        if (s[i][j] == 4)
                        {
                                HANDLE hOut;
                                hOut = GetStdHandle(STD_OUTPUT_HANDLE);
                                SetConsoleTextAttribute(hOut, BACKGROUND_RED |
BACKGROUND_GREEN | BACKGROUND_BLUE | BACKGROUND_INTENSITY | 7);
                                cout << " 4";
                        }
                }       cout << "\n";
        }

// Data output
        HANDLE hOut;
        hOut = GetStdHandle(STD_OUTPUT_HANDLE);
        SetConsoleTextAttribute(hOut, BACKGROUND_RED | BACKGROUND_GREEN |
BACKGROUND_BLUE | BACKGROUND_INTENSITY);

        cout << " Number of iterations: " << iteration << "\n";
        cout << " The number of individuals of Species 1: " << a << "\n";
        cout << " The number of individuals of Species 2: " << b << "\n";

// Calculating the states of the lattice sites in the next iteration
        for (int i = 0; i<n; i++)
        for (int j = 0; j<m; j++)

/* Calculating the state of the lattice site in the next iteration. The arguments of the
f1 function are the state of the central site and the states of sites from its
neighborhood. */
        s_temp[i][j] = f1(s[Torus_n(i)][Torus_m(j)],
                                s[Torus_n(i - 1)][Torus_m(j)],
                                s[Torus_n(i - 1)][Torus_m(j + 1)],
```

```
                                        s[Torus_n(i)][Torus_m(j + 1)],
                                        s[Torus_n(i + 1)][Torus_m(j)],
                                        s[Torus_n(i + 1)][Torus_m(j - 1)],
                                        s[Torus_n(i)][Torus_m(j - 1)]);

/* Transfer of new states of lattice sites to an array of current states of lattice
sites. */
            for (int i = 0; i<n; i++)
            for (int j = 0; j<m; j++)
            s[i][j] = s_temp[i][j];

            _getch(); // Pause
            iteration++; // Iteration counter
            system("cls"); // Clearing the screen
      }
      return 0;
}
```
